# *ZATELLITE:* a toolkit to visualize and manipulate human centromeres in live cells using synthetic zinc fingers

**DOI:** 10.64898/2026.08.04.742839

**Authors:** Nestor Saiz, Aleah Goldberg, Finnegan Clark, Guohong Tong, Manjunatha Kogenaru, Peter H Whitney, Osama Abdin, Fabio Alfieri, Philip M. Kim, Glennis A. Logsdon, Liam J. Holt, Jef D. Boeke, Marcus B. Noyes, Teresa Davoli, Timothee Lionnet

**Affiliations:** Institute for Systems Genetics, NYU Grossman School of Medicine, New York, NY, USA; Vanderbilt University School of Medicine, Nashville, TN, USA; Department of Molecular Genetics, University of Toronto, Toronto, Ontario, Canada; Department of Computer Science, University of Toronto, Toronto, Ontario, Canada; Donnelly Centre for Cellular and Biomolecular Research, University of Toronto, Toronto, Ontario, Canada; Department of Genetics, Epigenetics Institute, Perelman School of Medicine, University of Pennsylvania, Philadelphia, PA, USA; Department of Biochemistry and Molecular Pharmacology, NYU Grossman School of Medicine, New York, NY, USA; Department of Biomedical Engineering, NYU Tandon School of Engineering, Brooklyn, NY, USA; Department of Computational and Systems Biology, University of Pittsburgh School of Medicine, Pittsburgh, PA, USA; Department of Cell Biology, NYU Grossman School of Medicine, New York, NY, USA

## Abstract

Changes in the number of chromosomes or their spatial organization within the nucleus have critical consequences for cell fate. Yet capturing the karyotype or three-dimensional architecture of chromatin in living cells remains limited by the difficulty in labeling endogenous loci non-invasively. The most widely used tools rely on dCas9, a bulky protein whose persistent DNA binding interferes with DNA and RNA metabolism, causes DNA damage, and is hard to multiplex. We developed *ZATELLITE*, an AI-enabled tool to target endogenous repetitive sequences with fluorescently-tagged synthetic zinc fingers. *ZATELLITE* probes have key advantages over dCas9: they are smaller, easier to multiplex and, critically, they do not cause DNA damage or chromosomal abnormalities, even after long-term labeling. We generated a collection of *ZATELLITE* probes to label the centromeres of nearly all human chromosomes, enabling the capture of genome organization and karyotype alterations in real time in living cells. Finally, we show that *ZATELLITE* can be used for simultaneous labeling and epigenetic editing of centromeres. Thus, *ZATELLITE* is a non-toxic, versatile tool for genome visualization and manipulation.

## Introduction

The chromosomes of eukaryotic cells need to be tightly packed within the cell nucleus while also enabling gene expression. Genome organization is thus tightly regulated: chromosomes occupy distinct nuclear territories in interphase, with chromatin compartments that contain domains of highly associating chromatin^1^. This hierarchical organization must also be dynamic, to adapt to changing needs during the cell cycle and over the cell’s and organism’s lifespan. Thus, understanding the reciprocal relationship between genome organization and gene expression is fundamental for our understanding of cell physiology.

A wealth of studies using fluorescent in situ hybridization (FISH) and chromosome conformation capture techniques have established a 3D atlas of genome organization and chromatin interactions down to nucleotide resolution ^2–7^. However, our understanding of the temporal dimension of these interactions is hindered by limitations of existing tools to visualize genomic loci in living cells. The integration of operator arrays enables the imaging and tracking of multiple loci^8–11^ but is disruptive to the locus and labor intensive, as it requires editing of the genome of the cell of interest. The use of CRISPR/Cas9-derived tools^12^ overcomes the need for genome editing, reducing targeting to a gRNA design problem. However, despite ingenious improvements^13–18^, multiplexing remains challenging and the size of the dCas9-gRNA complex limits delivery. Moreover, dCas9 binding to DNA induces replication stress and chromosomal abnormalities^19–21^. Therefore, an alternative approach that overcomes these limitations would transform our ability to visualize the living genome.

Zinc fingers (ZFs), the most common DNA-binding domain in mammalian transcription factors, are small, modular and highly sequence-specific. While they have historically been difficult to design, we recently built a machine learning model, ZFDesign, that generates arrays of synthetic ZFs targeting any DNA sequence, enabling efficient transcription factor reprogramming and nuclease engineering^22^. We reasoned that fusions of fluorescent proteins (FPs) to designer ZFs would represent a compact tool to label genomic loci with minimal disruption to chromatin. Here we present *ZATELLITE* (Zinc finger Arrays Targeting Endogenous Loci to LIght up Tandem repetitive Elements), a toolkit that uses synthetic ZF arrays to label endogenous genomic loci. We validate *ZATELLITE* by targeting individual human centromeres, complex regions composed of long tandem repeats of α-satellite higher-order repeat (HOR) arrays. Centromeres are the site of attachment of the kinetochore and, as such, essential to maintain chromosome integrity and normal ploidy. However, because of their highly repetitive nature, they have traditionally been excluded from genomic analyses and our ability to study their biology has been limited. The recent sequencing of human centromeres finally makes it possible to capture the dynamics of these critical genomic elements^23,24^.

Here we show that labeling with *ZATELLITE* is comparable in quality to dCas9, but relies on probes that are 65-85% smaller and can be readily multiplexed. Critically, we demonstrate that, unlike dCas9, ZFs induce no DNA damage or chromosomal abnormalities when labeling centromeres. We have generated a collection of ZFs that uniquely label almost every human chromosome, enabling karyotyping of individual cells in vivo. Lastly, we demonstrate the versatility of these ZFs, which can be used not only to label, but also to manipulate genomic loci, enabling simultaneous visualization and epigenetic editing of individual centromeres. Thus, *ZATELLITE* provides a compact, safe and highly programmable alternative to dCas9 that can be used to visualize and manipulate genomic loci in living human cells.

## Results

### Synthetic fluorescent zinc finger arrays faithfully label endogenous repetitive loci in live cells

We have previously shown that synthetic zinc finger arrays can be easily designed to target endogenous loci using ZFDesign^22^. Here, we sought to target individual human centromeres, composed of long arrays of α-satellite HORs, which contain sequences unique to each chromosome, allowing specific targeting^23,24^. To test our approach, we selected 18bp target sequences found within the *DZ71* α-satellite HORs of the chr7 centromere as input for ZFDesign. We synthesized the coding sequences for ZF arrays predicted to bind to those sequences, fused them to a FP, and introduced the resulting plasmids into human cells. Binding of multiple units of the FP:ZF fusion to the tandem repeats should be visible as two fluorescent foci within the nuclei, marking the centromeres of the homologous chromosomes (Figure 1a). Accordingly, the vast majority of diploid human colonic epithelial cells (hCECs) stably infected with a fusion of the ZF to the green fluorescent protein mStayGold^25^ showed two nuclear foci (Figure 1b), whereas a clonal line containing an additional copy of chr7^19^ overwhelmingly showed three foci (Figure 1c), consistent with chr7 targeting.

**Figure 1.**
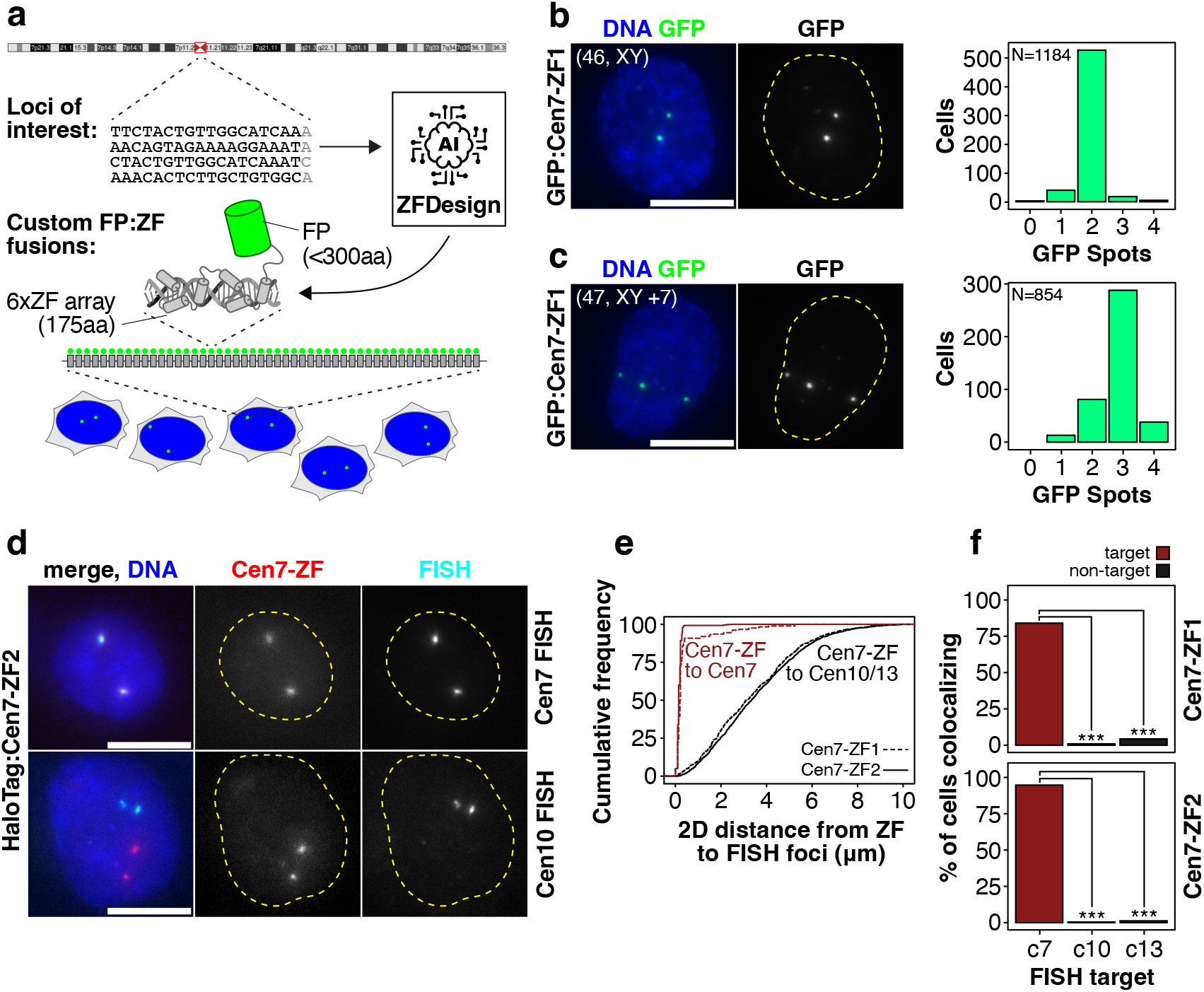
Synthetic fluorescent zinc finger arrays faithfully label endogenous repetitive loci in live cells. **a**. Diagram illustrating the generation of *ZATELLITE* zinc finger fusions to label genomic loci. Targets within a repetitive locus of interest are used to predict the sequence of ZF arrays that will recognize them using ZFDesign (Ichikawa et al., 2023). 18bp sequences targeted by the ZF array are indicated in black. The synthetic ZF array is fused to a fluorescent protein of choice and when introduced into cells, it binds the repetitive locus and appears as a fluorescent spot (or pair of spots). **b**. Representative images of a live diploid hCEC expressing an mStayGold:Cen7-ZF fusion, and the corresponding histogram indicating the number of fluorescent foci per nucleus across a population. **c**. Representative images of a diploid hCEC clone carrying an extra copy of chr7^19^ expressing the same mStayGold:Cen7-ZF fusion as in (b), with the corresponding histogram showing the distribution of foci per nucleus across a population. **d**. Representative images of HCT116 cells expressing a HaloTag:Cen7-ZF fusion labeled with JFX554, fixed and subjected to DNA FISH for cen7 (target) or cen10 (non-target control). **e**. Cumulative frequency distribution of distances between the HaloTag foci and the 2D nearest neighbor (NN) FISH focus, for either the target locus (dark red line) or non-target loci (cen10, cen13; black line) for two different ZF arrays targeting cen7, as indicated (and as shown in b-d). **f**. Fraction of HaloTag foci in (e) within 300nm of the nearest FISH spot (considered as co-localizing) for the target locus (cen7) or two non-target loci (cen10, cen13). *** = p<0.001 (Fisher’s Exact test). Scale bars = 10μm.

To validate that our synthetic FP:ZF fusions bind on target, we generated HaloTag:ZF fusions for two different Cen7-ZFs, stably integrated them into HCT116 cells using the PiggyBac system^26^ and then performed DNA FISH on them after labeling HaloTag with a suitable fluorophore. We observed co-localization of HaloTag:ZF foci and FISH foci for the centromere of chr7 (Cen7), but not with FISH foci for chr10 (Figure 1d). HaloTag:ZF foci had a median distance of <200nm to the nearest DNA FISH spot for Cen7, but a median distance of >3μm to a non-target locus (Figure 1e). To determine co-localization, we used distances obtained from co-labeling of a single locus by DNA FISH (see methods, Figure S1). While 98% of foci for Cen7-ZF1 and 88% of foci for Cen7-ZF2 were within 300nm of the nearest FISH focus for the target locus, only 1% or 2.9% were within 300nm of the nearest FISH focus for a non-target locus (cen10 or cen13, respectively) (Figure 1f), indicating high target specificity.

Thus, we can easily generate synthetic FP:ZF fusions that specifically bind to their target locus, enabling chromosome copy number capture in living cells. We call this tool *ZATELLITE*.

### ZATELLITE does not cause DNA damage or aneuploidy

We next wanted to compare *ZATELLITE* to dCas9, the current state of the art for labeling loci in living cells without genome editing. We used hCECs, which we have previously successfully used to label centromeres using dCas9^19^. We generated a diploid hCEC line stably expressing an mStayGold:dCas9 fusion by lentiviral infection and FACS-sorted cells to approximately match the fluorescence intensity of hCECs infected in parallel with a mStayGold:Cen7-ZF fusion (Figure S2a-b). We subsequently infected the resulting dCas9:mStayGold hCEC line with sgRNAs targeting the centromeres of either chr6, 7 or 18, and, in parallel, we infected the parental hCEC line with mStayGold:ZF fusions targeting the same centromeres (Figure 2a, S2c). After 5 days of positive selection, we confirmed successful locus labeling (Figures 2a, S2c), and established that both experimental groups had a comparable fraction of cells expressing mStayGold and nuclear fluorescence levels (Figure S2d-e). As expected, all cells showed predominantly two foci for each chromosome with either dCas9 or *ZATELLITE* (Figure S2f). Interestingly, our hCEC line often showed three foci for chr18, detected by both dCas9 or ZFs (Figure S2f), suggesting the presence of a chr18 triploid subpopulation. Indeed, single-cell DNA sequencing confirmed the presence of cells carrying an additional copy of chr18, but not of chr6 or 7, in our parental hCEC line (Figure S2g), further supporting the ability of *ZATELLITE* to detect karyotypic alterations in living cells.

**Figure 2.**
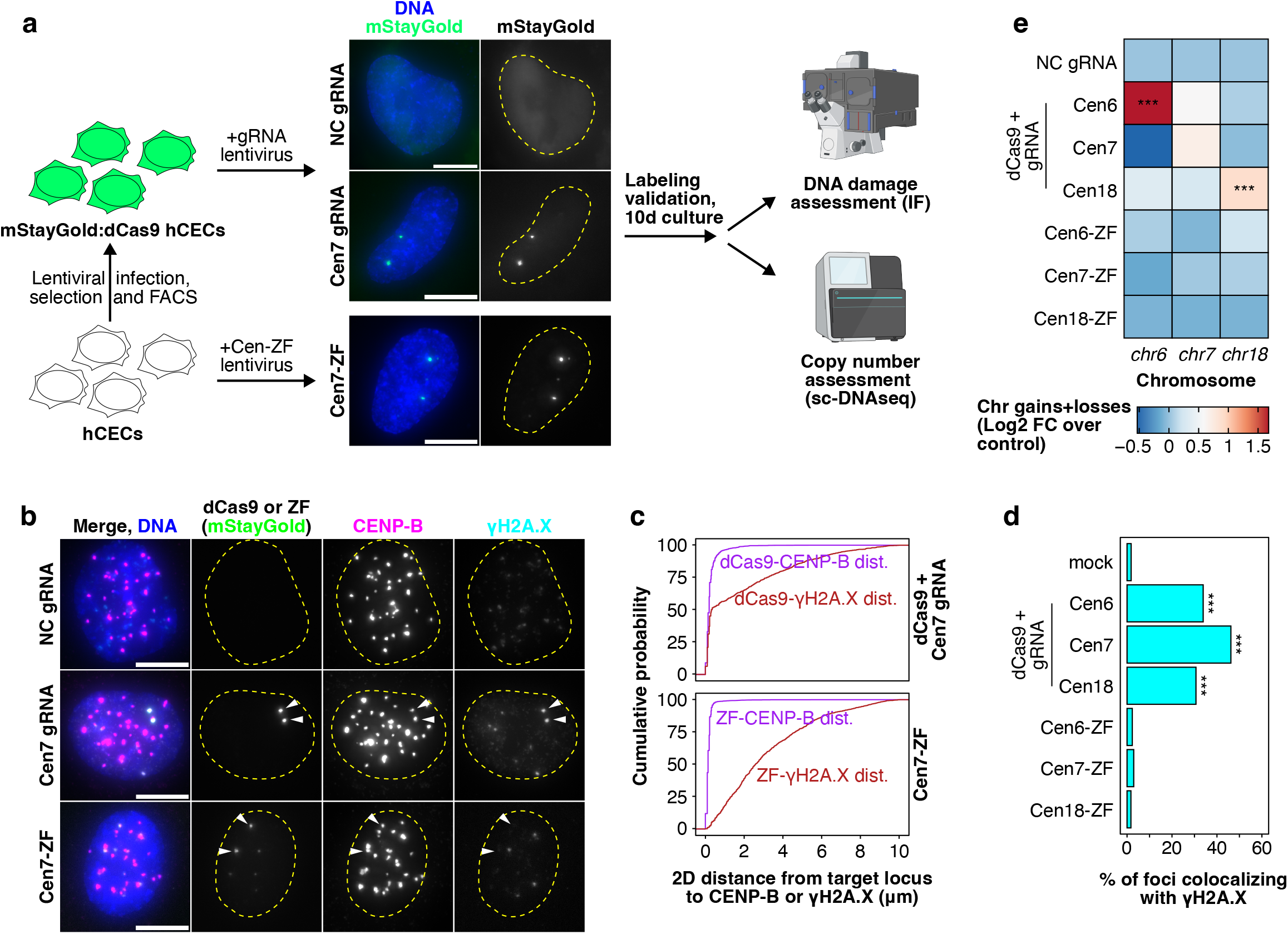
*ZATELLITE* does not cause aneuploidy or DNA damage. **a**. Diploid hCECs were first stably infected with a mStayGold:dCas9 fusion and subsequently infected with an sgRNA targeting cen7, as indicated. In parallel, unlabeled hCECs were infected with mStayGold:Cen7-ZF fusions. Images show hCECs 5-6 days post-infection displaying two fluorescent foci, corresponding to the centromere of chr7, bound by either dCas9 or the Cen7-ZF. Cells were allowed to proliferate for 10 days post-infection before performing immunofluorescence or scDNA-seq. **b**. Representative images of hCECs from the same population, fixed 10 days post-infection and stained for γH2A.X and CENP-B, a centromeric marker. Arrowheads indicate the position of the mStayGold (dCas9-sgRNA or ZF) foci across channels. **c**. Cumulative frequency distributions of the 2D nearest-neighbor (NN) distance between mStayGold foci and centromeres (marked by CENP-B, magenta) or γH2A.X foci (dark red). Distances below 300nm indicate co-localization. **d**. Fraction of mStayGold foci within 300nm (co-localizing) of the nearest γH2A.X foci in each experimental group across the population. **e**. Heatmap showing whole-chromosome gains/losses of chr6, chr7 or chr18 for cells infected with either sgRNAs or mStayGold:ZFs and cultured for 10 days post-infection, as indicated in (a). Fold-change (FC) relative to control is showing as log2. In all panels, *** = p<0.001 (Fisher’s Exact test). Scale bars = 10μm.

dCas9 binding to telomeres induces replication stress and DNA damage^20^, and dCas9 targeting human centromeres generates aneuploidy^21^. These observations suggest that the binding of imaging probes might lead to genomic instability. We first sought to establish whether *ZATELLITE* probe binding to centromeric loci induces DNA damage. We kept hCECs labeled with either dCas9-sgRNAs or ZFs growing for 10 days post-infection and performed immunofluorescence for the histone variant γH2A.X, a marker of double strand breaks (DSBs), as well as CENP-B, a pan-centromeric marker (Figure 2a). As expected, mStayGold (fused to either dCas9 or the ZFs) formed nuclear foci co-localized with CENP-B in all groups, confirming successful targeting of centromeres (co-localization frequency >75%; Figure 2b-c, S2h-i). Strikingly, we observed that in cells labeled with dCas9, 35-50% of the mStayGold:dCas9 foci co-localized with a bright γH2A.X focus (Figure 2b-d, S2h-i). In contrast, in cells expressing mStayGold:ZF fusions, <4% probe spots co-localized with a γH2A.X focus, not significantly different from the frequency of random foci in control groups co-localizing with γH2A.X or the apparent co-localization frequency between unrelated loci (Figure 1e, 2b-d, S2h-i). Assigning individual cells to their cell cycle stage demonstrated that cells bearing γH2A.X foci at the target locus were predominantly in late S and G2 (Figure S2j), suggesting that DNA damage accumulates as a result of DNA replication stress, as previously shown^20^.

We then asked whether the extensive levels of DNA damage we observed at centromeres could lead to genomic instabilities. To this end, we analyzed chromosome copy number at 10 days post-infection using single-cell DNA sequencing. Centromeric labeling with dCas9-sgRNA caused an increase in gains or losses of the arms of the targeted chromosome for all three loci we tested (chr6, 7 and 18) (Figure 2e, S2k, Tables S1-3), whereas in cells where the same centromere was targeted with ZFs, we only observed background levels of gains/losses, not significantly different from those of cells infected with a non-coding sgRNA (NC gRNA) (Figure 2e, S2k, Tables S1-3). We observed an increased rate of gains/losses at the whole chr7 level in cells infected with gRNA targeting chr7, though not statistically significant compared to the control (Figure 2e, Table S2). This is likely driven by the non-significant increase in chr7p gain observed when analyzing arm-level events (Figure S2k, Table S3), possibly due to under-sampling in this population. This effect was specific to the target locus, with non-targeted centromeres showing no copy number alteration above baseline with either labeling method (Figure 2e, S2k).

In summary, labeling using *ZATELLITE* is comparable in quality to labeling with dCas9, without causing aneuploidy or build up of DSBs at the centromeres, overcoming a significant limitation of dCas9-based methods for locus labeling.

### A collection of synthetic zinc finger arrays to label nearly all human centromeres

Human centromeres have complex, highly repetitive sequences that are largely unique to each chromosome^27^. Having established synthetic ZFs as a tool to specifically label the centromeres of chr6, 7 and 18, we sought to generate a panel of *ZATELLITE* probes to uniquely identify all centromeres in live human cells. We picked target sequences that were specific to each chromosome from a previously curated list^19^ and established they are conserved at the population level using a collection of 2,110 diverse centromeres from 65 individuals assembled by the Human Genome Structural Variation Consortium (HGSVC)^28^ (Figure 3a, S3a, Table S4). Target sequences for two pairs of the acrocentric chromosomes (chr13/21 and 14/22) were found in both members of each pair, consistent with their high degree of homology^23,24^. We then used ZFDesign to generate the sequences of ZFs predicted to bind each target sequence (as in Figure 1a). We screened a library of 224 candidate designs targeting all 24 human centromeres and selected for further analysis those that generated pairs of distinct fluorescent foci when transiently transfected into human cells (35/224), consistent with cell ploidy.

**Figure 3.**
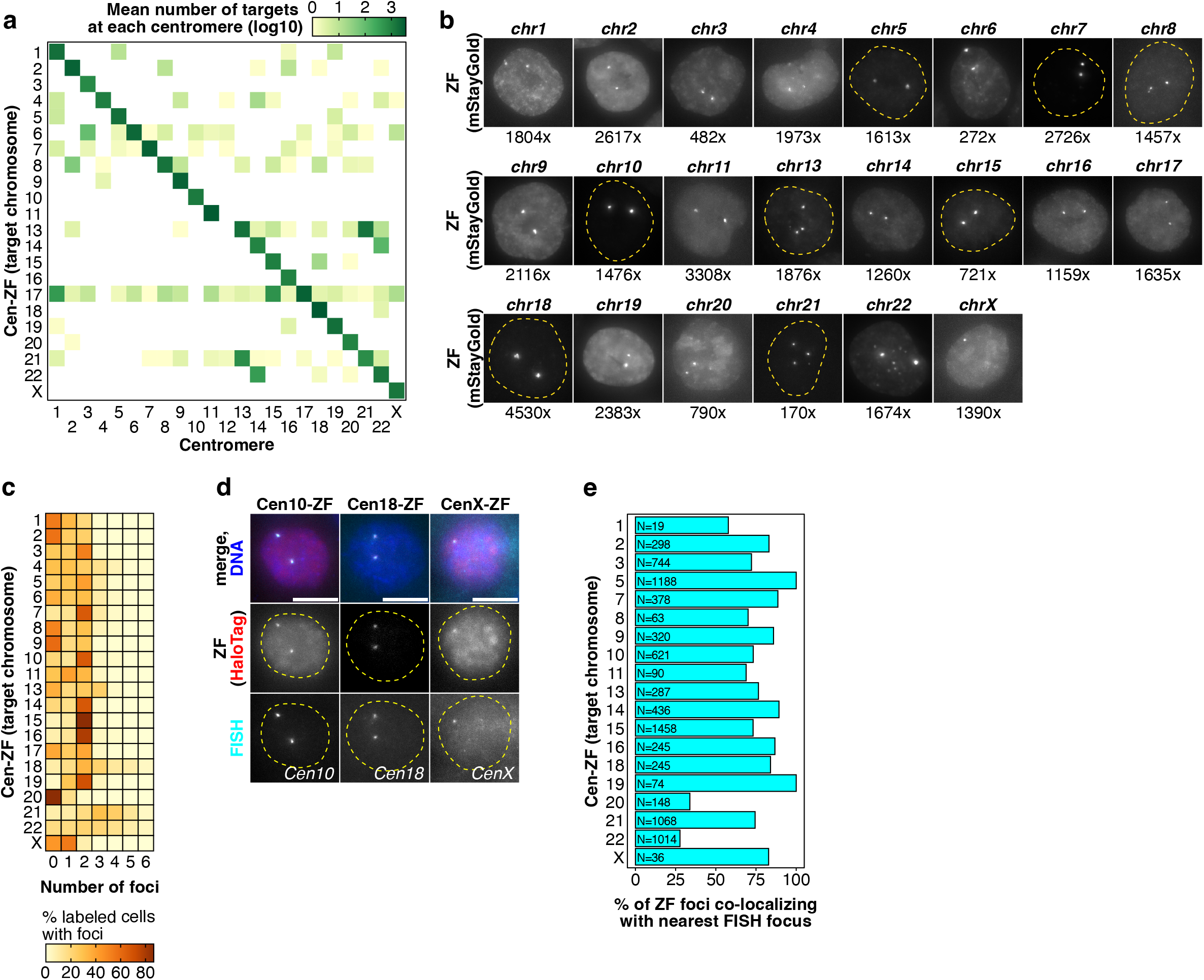
A collection of designer zinc finger arrays to label nearly all human centromeres. **a.** Heat map showing the mean number of targets (shown as log10) found for each Cen-ZF design in a set of 2110 human centromeres from a cohort of 65 individuals^28^. Individual values are shown in Figure S3a. **b**. Representative images of live HCT116 cells stably expressing mStayGold:Cen-ZF fusions targeting unique sequences within the centromeres of each of the 24 human chromosomes. The chromosome labeled is indicated over each image. The number of binding sites for each probe in the T2T genome is shown beneath each image. **c**. Heat map showing the number of foci per nucleus for each of the Cen-ZF fusions shown in a. Color-coding indicates the % of mStayGold+ nuclei with the indicated number of foci in each line. **d**. Representative images of HCT116 stably expressing HaloTag:ZF fusions targeting the centromeres indicated after being fixed and subject to DNA FISH for their target locus. HaloTag, labeled with JFX554, indicates the location of the *ZATELLITE* probe. The target loci (FISH) are marked by fluorescently labeled oligonucleoutides. **e.** Bar chart showing the % of Cen-ZF foci co-localizing with the nearest FISH focus for each probe in images like those shown in c. N indicates number of foci analyzed. Scale bars = 10μm.

We then used PiggyBac to integrate the selected ZF designs fused to mStayGold into HCT116 cells and generate polyclonal cell lines stably expressing these probes. Nearly all designs produced two distinct foci in most cells (34/35), reporting the location of each centromere pair for somatic chromosomes and of the single copy of the X chromosome in these cells (Figure 3b). We found a degree of variability in the labeling sensitivity across designs, resulting in variable number of foci (Figure 3c, S3b, Table S5) and signal to background ratio (SBR) (Figure S3c). We attribute these differences to a combination of noise in the image acquisition, spot detection and classification (see Methods), the variable affinities of each design for its target, the variable number of target sequences, or variable expression levels in polyclonal cell lines.

To validate probe specificity, we generated cell lines stably expressing fusions of each probe with HaloTag, which can be labeled with a spectrally diverse set of bright organic dyes, and assayed the co-localization of each probe with its target locus by DNA FISH (as in Figure 1d-e). We selected 15 probes targeting non-acrocentric chromosomes from our library for testing (Figure 3d, S3d), and confirmed that all mark the target locus based on co-localization between the ZF and the DNA FISH foci (Figure 3d-e, S3d).

Chr13 and 14 are acrocentric chromosomes whose centromeres share high sequence homology with chr21 and 22, respectively.^23,24^. As a consequence, CRISPR/dCas9 tools have so far failed to distinguish chromosomes in these pairs^21^ and FISH probes targeting the centromere of only one of the chromosomes are not available, to our knowledge. Consistently, the target sequences for our ZF designs were found in both loci for each pair (Figure 3a, S3a). As expected due to this homology, probes targeting the centromeres of chr13, chr21 and chr22 often showed more than two foci (Figure 3b, S3d-f). Due to the limitations of existing reagents, we validated acrocentric centromere ZF designs using commercially available DNA FISH probes targeting either the cen13/21 pair or the cen14/22 pair (Figure S3d). The Cen22-ZF probe often generates four spots per cell which colocalize with the cen14/22 FISH probes, demonstrating the probe labels both chromosomes 14 and 22. On the other hand, the probe for cen14 yields two distinct foci that co-localize with two of the spots labeled by the cen14/22 FISH probe (Figure S3d), suggesting that it can be used to resolve these loci (Figure S3e). Unfortunately, we were not able to distinguish cen13 from cen21 using our current probes (Figure S3f), perhaps reflecting a higher degree of homology between these two loci.

Nonetheless, our current validated probe set demonstrates the potential of this approach to readily generate probes to specifically and safely label all human centromeres.

### ZATELLITE enables direct, multi-color labeling of endogenous loci

Labeling multiple loci in distinct colors in the same cell with dCas9 is difficult: since the fluorescent label (dCas9:FP) is decoupled from the target specificity (gRNA), one needs to use complex aptamer-based systems^13,16,29^. In contrast, ZFs targeting different loci can be readily fused to spectrally distinct fluorescent proteins to label multiple loci at once. To demonstrate this capability, we labeled two or three Cen-ZFs with different FPs and integrated them into HCT116 cells using PiggyBac. Pairs of foci for each of the centromeres targeted could be easily identified in live cells after drug selection (Figure 4a, b). Human acrocentric chromosomes (13, 14, 15, 21, 22) carry the nucleolar organizer regions (NORs), containing rRNA genes, around which nucleoli are formed. We thus assessed the localization of some of the acrocentric centromeres relative to the nucleolus, identified by staining of nucleophosmin (NPM1) on 2D projections. As expected, the centromeres of chr14, 15 and 21 localized inside or on the edge of the nucleolus in 82-96% of cases (Figure 4c, d). By contrast, the centromeres of chr7 or chr10 were only found in the same regions in 63-64% of cases, showing comparable distribution between all three compartments (Figure 4c, d).

**Figure 4.**
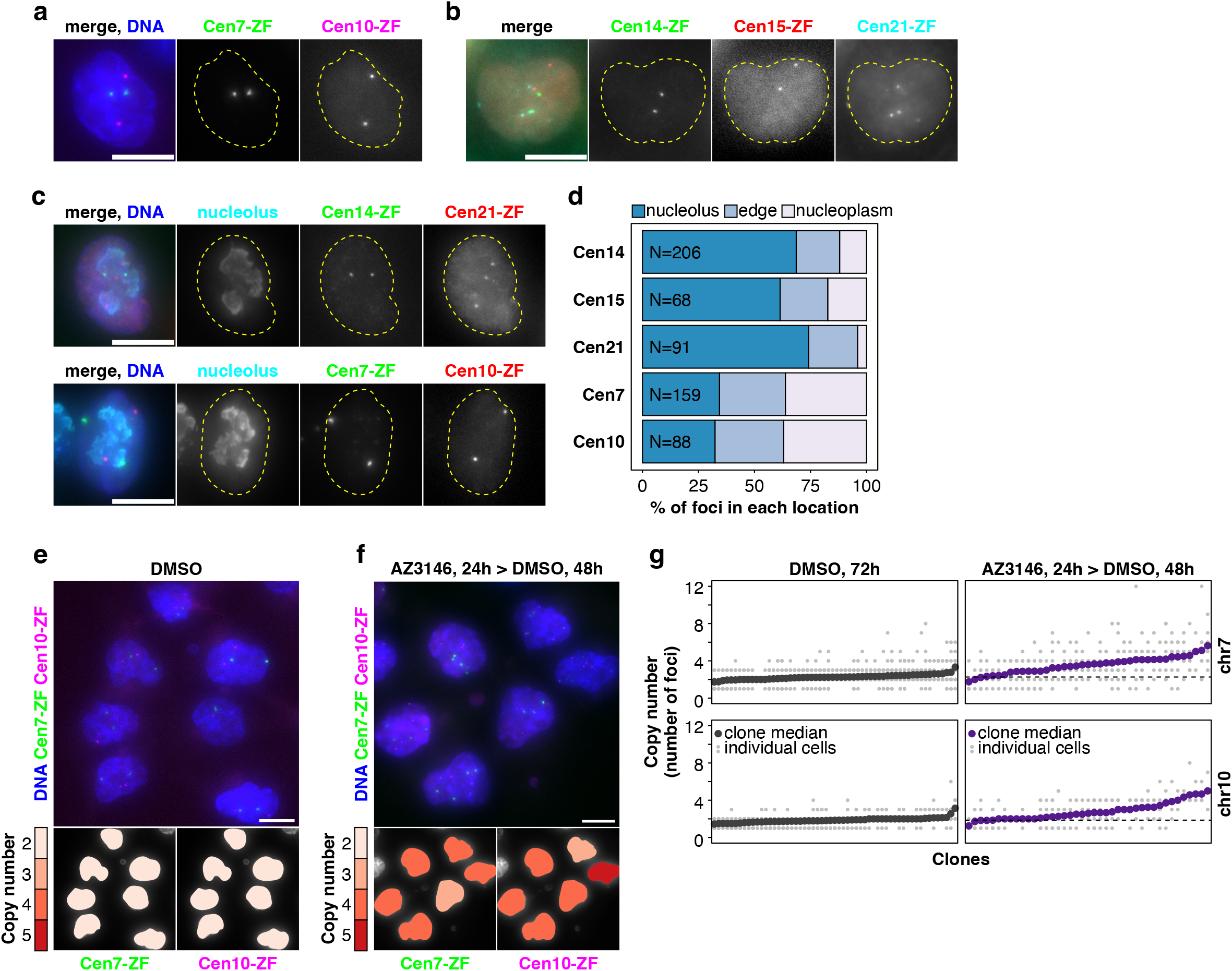
*ZATELLITE* enables direct, multi-color labeling of endogenous loci. **a.** Representative images of live HCT116 stably expressing Cen7-ZF fused to mStayGold and Cen10-ZF fused to mScarlet3-H^51^. **b.** Representative images of live HCT116 stably expressing Cen14-ZF fused to mStayGold, Cen15-ZF fused to mScarlet3-H and Cen21-ZF fused to HaloTag and labeled with JF657. In both a and b DNA is stained with Hoechst. **c.** Representative images of fixed HCT116 from the same lines as in a and b, stained for nucleophosmin (NPM1) to label the nucleolus. Images are 2D maximum intensity projections of 9μm stacks. **d.** Stacked bar plot showing the fraction of Cen-ZF foci in each of the indicated compartments in 2D projections as shown in c: nucleolus (overlapping with NMP1 staining), nucleoplasm (non-overlapping), or edge of the nucleolus (foci at the interface between the NPM1 compartment and the nucleoplasm). N indicates the number of nuclei analyzed for each locus. **e, f.** Representative images of clones of HCT116 cells from the line shown in a, grown in 0.1% DMSO for 72h (e) or in 5μM AZ3146 (24h), then in 0.1% DMSO (48h) before fixing. Nuclear masks on the right panels are color coded for the number of foci in each nucleus for each locus, as indicated in the legend below. **g.** Dot plot indicating the copy number in clones like those shown in e and f, treated as indicated. Each point in the X axis represents one clone: bold dots show the average copy number of each clone, gray dots behind them show the copy number of each cell in that clone. Dashed lines indicate the average copy number for control clones (DMSO) for each chromosome. Scale bars = 10μm.

Acquisition of aneuploidy is a hallmark of cancer cells, but its contribution to cancer progression is not fully understood^30^. In principle, labeling centromeres in living cells enables monitoring copy number longitudinally or tracking the fate of individual aneuploid cells within a population. To test whether *ZATELLITE* could be applied in this context, we induced aneuploidy in HCT116 cells expressing both Cen7-ZF and Cen10-ZF probes using an established approach. We inhibited MPS1 kinase, a regulator of the mitotic spindle assembly checkpoint, using 5μΜ AZ3146^31^ for 24h, followed by 48h without drug, and monitored foci counts in distinct clones. As expected, clones of control cells overwhelmingly retained two foci for both ZF probes, indicative of a diploid karyotype (Figure 4e), whereas clones of cells treated with the drug frequently showed 3-4 copies of each chromosome, with extreme cases having up to 8-12 copies (Figure 4f, g), confirming the expected widespread chromosomal gains for both chr7 and chr10. These data demonstrate that *ZATELLITE* is a powerful tool to monitor copy number alterations with single-cell resolution, in living cells.

### Simultaneous labeling and epigenetic editing of human centromeres in live cells using synthetic zinc fingers

Lastly, we sought to extend the use of *ZATELLITE* beyond live locus labeling. We hypothesized that target-bound ZF fusions may serve as a homing element to which another protein complex can be inducibly recruited. To that end, we devised the following two-component system (Figure 5a): a *ZATELLITE* probe fused to ABI1 (ABI1:Cen7-ZF:mStayGold), constitutively labels the target locus, while a second chimeric element, a catalytic domain (CD) fused to both HaloTag and PYR1 (NLS-HaloTag:CD:PYR1), is only recruited to the target locus upon the addition of the small molecule Mandipropamid (Mandi) which mediates ABI1-PYR1 dimerization^32^. The fast and reversible kinetics of Mandi provide fine temporal control of the recruitment, while HaloTag enables visualization of the chimeric CD to confirm successful recruitment (Figure 5b).

**Figure 5.**
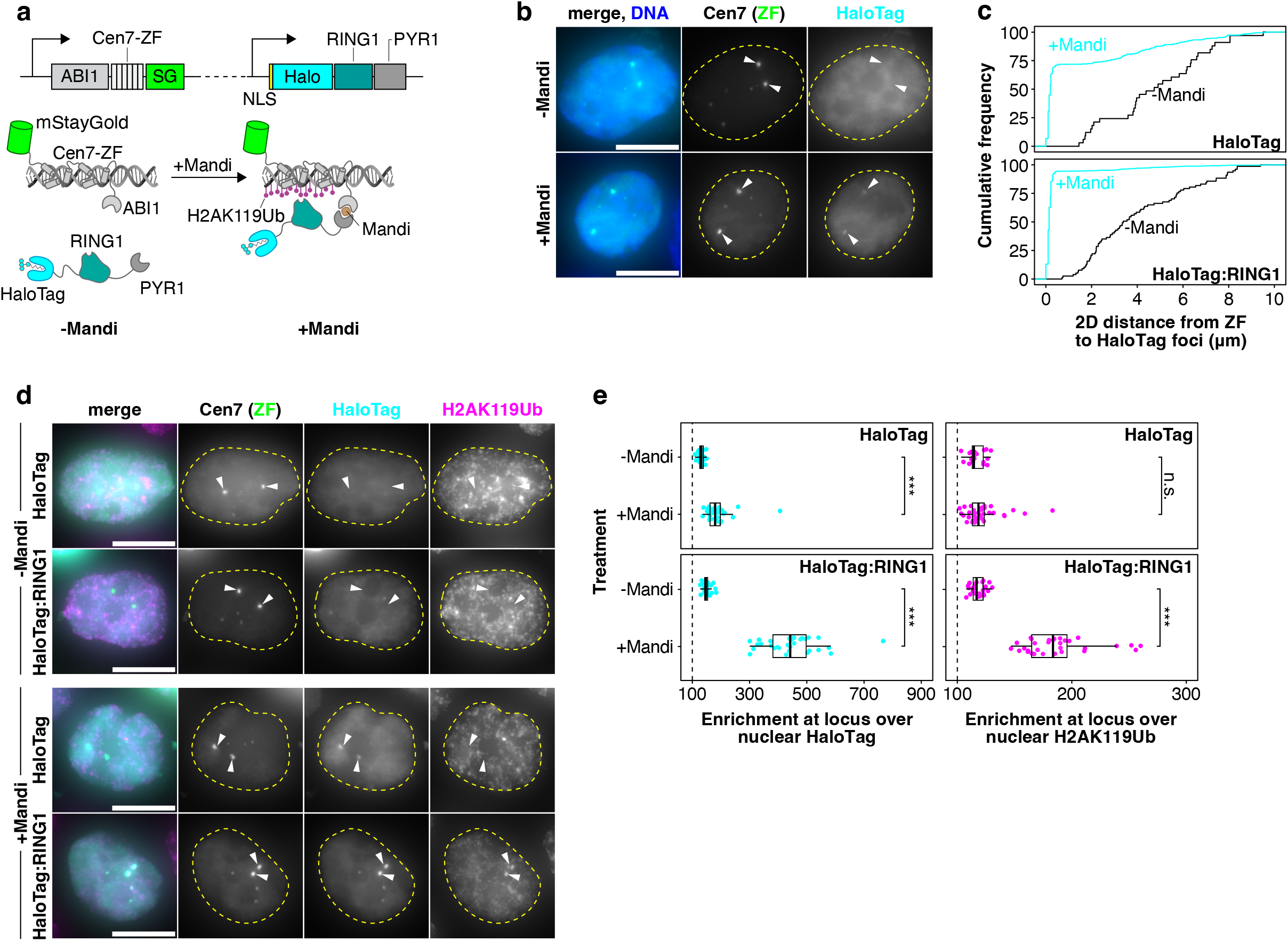
Simultaneous labeling and epigenetic editing of human centromeres in live cells using synthetic zinc fingers. **a.** Diagram illustrating the two-component system to label and edit genomic loci in living cells. A constitutive promoter drives the expression of the *ZATELLITE* fluorescent Cen7-ZF fused to ABI1, one of the interaction domains to achieve chemically induced proximity (CIP). The second component is a fusion of HaloTag, RING1 and PYR1 (“HaloTag:RING1” in the rest of the panels). PYR1 dimerizes with ABI1 in the presence of Mandipropamid (Mandi), thus recruiting HaloTag:RING1 to the labeled locus. RING1 can be replaced with any other catalytic domain, or an “empty” fusion can be used instead (HaloTag in the rest of the panels). **b**. Representative images of HCT116 cells transfected with the empty version of the construct shown in (a) expressing only the HaloTag:PYR1 fusion. Treatment with Mandi induces recruitment of HaloTag:PYR1 to the chr7 centromere, labeled by the ZF:mStayGold fusion. **c**. Cumulative frequency distributions of the 2D nearest-neighbor (NN) distances between mStayGold (ZF) foci and HaloTag foci in control cells treated with 0.1% DMSO (“-Mandi”, black curves) or in cells treated with 5μM Mandi (cyan curves). Addition of Mandi results in co-localization of most ZF foci with HaloTag foci. **d**. Representative images of HCT116 cells transfected with either the emptyversion of the construct shown in (a) (HaloTag:PYR1) or the construct expressing RING1 (HaloTag:RING1:PYR1), treated with either 0.1% DMSO (“-Mandi”) or 5μM Mandi. Arrowheads indicate the position of the Cen7, as labeled by mStayGold:Cen7-ZF. Recruitment of HaloTag:RING1:PYR1 to the target locus results in deposition of H2AK119Ub at the chr7 centromere. **e**. Box plots showing the enrichment of either HaloTag or H2AK119Ub at the locus labeled by the Cen7-ZF in each of the conditions shown, for either the HaloTag or the HaloTag:RING1 construct. Enrichment for each protein is calculated as the intensity at the spot divided by the median intensity of the nucleus and represented as log2. Each dot indicates the average enrichment in all cells in one image (∼15 cells/image). *** = p < 0.001 (one-way ANOVA). n.s = not significant. Scale bars = 10μm.

As a proof-of-principle, we designed a chimera carrying a nuclear HaloTag:RING1:PYR1 fusion, as well as a control chimera consisting only of HaloTagged PYR1 (NLS-HaloTag:PYR1). RING1 is a component of the Polycomb Repressive Complex (PRC) that mediates the ubiquitination of H2AK119. Upon transfection of a plasmid co-expressing the chimera and mStayGold:Cen7-ZF:ABI1 into HCT116 cells, both HaloTag chimeras localized to the nucleus of cells, while the ZF probe formed two fluorescent nuclear foci, as expected (Figure 5b). Addition of 5μM Mandi caused the appearance of HaloTag foci co-localizing with the ZF foci, consistent with recruitment of the chimeras to Cen7 (Figure 5b, c). To probe the biological function of the recruited chimera, we treated cells with 5μM Mandi for 6h before fixing the cells and assessing the presence of H2AK119Ub by immunofluorescence (Figure 5d). We observed that recruitment of RING1 to the target locus resulted in a near 2-fold accumulation of H2AK119Ub at the target locus over its average nuclear level (Figure 5d, e). By contrast, in cells not treated with Mandi or in cells where we recruited the control chimera (HaloTag:PYR1 fusion) to Cen7, there was no significant accumulation of H2AK119Ub at the locus (Figure 5d, e).

These data demonstrate that synthetic ZFs provide a platform to not only readily label genomic loci in living cells, but to alter their epigenetic state at will. Their small size and potential for engineering make them a powerful and physiologically safe alternative to dCas9-based methods.

## Discussion

In this study we introduce *ZATELLITE*, a toolkit to label and manipulate multiple repetitive genomic loci in living mammalian cells using synthetic fluorescent ZF arrays. As part of its development, we have generated a collection of validated ZFs to uniquely label the centromeres of nearly all human chromosomes. Our approach leverages ZFDesign^22^ to generate candidate ZF arrays that, once screened, demonstrate high specificity for their target sequence. *ZATELLITE* probes are small, facilitating their delivery, versatile, in that they can be readily fused to any protein domain, and modular, allowing fusion to functionally diverse domains to carry out complex processes, such as simultaneous labeling and epigenetic modification.

Current methods to fluorescently tag loci without genome editing are powerful but present limitations: CRISPR/dCas9-based methods are limited when it comes to delivery, due to the large size of dCas9 (∼1400aa, ∼500aa for CasMINI^33^), and multiplexing. Because the fluorophore is fused to dCas9, while target specificity is supplied by a gRNA, every locus bound by a gRNA is necessarily labeled in the same color. Assigning different colors to different loci thus requires either orthogonal Cas proteins or engineered sgRNA scaffolds carrying distinct fluorophore-recruiting aptamers, approaches that are cumbersome and cap the number of loci that can be resolved at once^13,29^. Transcription activator-like effector (TALE) domains can also be used for labeling^34,35^. Like ZFs, TALEs encode sequence specificity in the protein, but are also limited by their size (>600aa for an 18bp target) and their highly repetitive nature. By contrast, a 6xZF array targeting an 18bp target is 175aa long. Critically, we show that, unlike dCas9, labeling using ZFs does not cause DNA damage or loss of chromosomal arms. We speculate that this difference comes from the more transient interaction kinetics of ZFs with DNA relative to dCas9. Whereas canonical dCas9-gRNA complexes have an estimated residence time of >3h^36^, *ZATELLITE* probes dissociate from their targets on the order of seconds (not shown). Persistent binding of hundreds of dCas9-gRNA complexes at centromeres, with the associated unwinding of the DNA helix, would conceivably interfere with the DNA replication machinery, accounting for the reported replication stress^20^, and result in chromosomal arm losses^19,21^, (this study). Although DNA lesions might be less frequent when targeting non-centromeric regions, it remains unknown which locus features underlie sensitivity to dCas9 targeting, justifying the need for an alternative method.

The availability of ZFDesign, a novel tool to predict ZF-DNA interactions^22^, and of the complete sequence of human centromeres released by the T2T consortium^23,24^ allowed us to engineer a collection of ZF probes targeting nearly all human chromosomes. These probes show remarkable specificity — of 24 designs (targeting 19 centromeres) tested by DNA FISH, 22 showed specificity for their target locus. We have yet to generate or validate designs for 5 centromeres (chr4, chr6, chr12, crh17 and chrY), but our success so far suggests these are within reach through more screening. Crucially, *ZATELLITE* is not limited to targeting centromeres: we have generated probes to label other repetitive elements, including telomeres and minisatellites (not shown), which extend the application of this tool to any repetitive element in the genome.

A current limitation of *ZATELLITE* is a moderate efficiency of candidate designs, which requires screening multiple probes per locus. Continued improvements to the design and screening pipeline should circumvent this challenge. Likewise, while validated probes are highly specific, they show variable signal-to-background ratio (SBR) and sensitivity, indicating that probe performance depends not only on target copy number but also on design features that remain to be optimized (Figure S3c). Importantly, probes currently expressed from randomly integrated constructs in polyclonal populations already generate foci that are sufficiently robust for routine live-cell imaging applications, underscoring the practical usability of the platform (Figure 4). Additional gains in signal quality should be achievable through single-copy integration, signal amplification strategies, and rational tuning of probe affinity^37^.

Here we demonstrate detection of foci with as little as 170 target sites. This number is unlikely to represent the absolute detection floor, since single mRNA molecules decorated with six binding proteins can be tracked in living cells^38^. Given the small size of zinc fingers, it should be feasible to express a limited set of optimized probes tiling a non-repetitive locus, opening the way to safe labeling of arbitrary endogenous genomic sites in living cells without genetic engineering through future iterations of *ZATELLITE*.

Lastly, we demonstrate the versatility of *ZATELLITE* by extending the use of ZF probes beyond labeling, to epigenetic editing of a target locus with high temporal control. Our ability to manipulate centromeres has so far been mostly limited to the use of artificial centromeres^39,40^, to pan-centromeric studies^41^ or to the use of dCas9^19,21^, with its associated side effects. Our collection of specific centromeric ZFs opens the door to more selective studies targeting individual, or groups of, chromosomes of interest, without the risk of damage to the subject of study.

In conclusion, synthetic ZFs represent a versatile and safer alternative to dCas9-based methods for genome visualization and manipulation that overcomes their main limitations. *ZATELLITE* probes targeting individual human centromeres are a powerful tool to visualize and manipulate these critical genomic regions that have so far remained largely inaccessible. We expect further development of this technology to greatly expand the synthetic biology toolkit and enable large-scale chromatin and even chromosome level manipulations.

## Methods

Manufacturers and catalog numbers for all relevant reagents have been compiled in Tables S6 and S7. These details are thus omitted from the text.

### ZF probe selection and design

We mainly generated synthetic arrays of 6xZFs using ZFDesign^22^ with 19bp target DNA sequences as input (Fig. 1a). ZFDesign was developed to generate arrays composed of ZF pairs separated by a base-skipping linker. To design arrays of 6 contiguous ZFs, we used a modified sampling procedure to generate longer arrays. During array design, we computed amino acid probabilities for ZFs within the array using context from both neighbors independently. The minimum probability for each amino acid across the two contexts was used when sampling residues at each position. The result of this procedure was a longer array of mutually compatible ZFs (6 in our case, with a length of 522bp (174aa)). For three centromeres (chr7, chr18, chrX) we also generated arrays of 8xZFs using 28bp targets as input. These arrays were composed of four compatible pairs, separated by a base-skipping linker (729bp, or 243aa).

Candidate 19bp target sequences were chosen from a previously curated list of 20-bp sequences found in the live higher-order repeats (HORs) of human centromeric α-satellite repeats^19^. 28bp targets were chosen from an in-house generated list of 35bp sequences. Selected 19bp targets met the following criteria: (1) chromosome and centromere specificity score of 0.85 or higher (most with scores 0.99-1) and (2) no more than 3 instances of any of the following triplets: NNG, CGC, TGC, AGC, TTC, CTC or TTA. For each centromere, we generated 6-24 ZF arrays for putative targets found in either strand, for a total of 224 ZF arrays.

### Plasmid construction and screening

Predicted DNA sequences encoding ZF arrays were codon-optimized using the IDT codon optimization tool (http://idtdna.com/codonopt) and a *Homo sapiens* codon usage table. Codon-optimized sequences were flanked by constant 15bp homology arms and purchased as gene fragments from IDT (either as gBlocks^TM^ or eBlocks^TM^). Synthetic gene fragments were directly assembled into linearized plasmid vectors using NEBuilder HiFi DNA Assembly Master Mix following manufacturer instructions (https://dx.doi.org/10.17504/protocols.io.bfhrjj56). Destination vectors were either pcDNA3.1 NT-GFP-TOPO derivatives or PiggyBac vectors that were linearized by PCR using Q5 High Fidelity Master Mix or PrimeSTAR GXL Premix. PCR products were purified using Zymoclean Gel DNA Recovery Kit. Assembly products were transformed into NEB 5-alpha competent *E coli* using the manufacturer’s protocol (https://dx.doi.org/10.17504/protocols.io.bddti26n) and grown on LB medium under ampicillin selection. When possible, volumes and DNA amounts were scaled down to maximize reagents. Colonies of transformed *E coli* were screened by colony PCR and liquid cultures from candidate samples grown for plasmid extraction using the QIAprep Spin Miniprep Kit. Plasmid sequences were verified by whole-plasmid sequencing (Plasmidsaurus). The same workflow was used to generate and clone other DNA fragments, such as mStayGold or HaloTag®^42^.

GFP:ZF designs cloned into pTOPO and driven by a CMV promoter were initially screened by transient transfection into U2-OS or HCT116 cells and live imaging 24-48h later. ZF designs showing the expected number of nuclear foci in transfected cells were subcloned into a PiggyBac vector for stable integration and further characterization. In the PiggyBac vector, ZFs were fused with either mStayGold or HaloTag and driven by an EF1α promoter.

Barcoded lentiviral vectors for mStayGold:dCas9 and mStayGold:Cen-ZF transduction were modified from the pHAGE-3xmScarlet-dCas9 vector used in^19^ by Gibson cloning. Lentiviral vectors expressing sgRNAs were previously used in^19^. Constructs used for ZF recruitment experiments were derived from the Watermelon lentivirus^43^. The open reading frames in the Watermelon vector were modified using synthetic gene fragments or PCR products combined by Gibson assembly (NEBuilder HiFi) and then subcloned into a PiggyBac vector.

### Computational quantification of Cen-ZF target and off-target sequences

To estimate the abundance of candidate binding sites for each Cen-ZF design, target sequences were searched against the T2T-CHM13-derived centromeric HOR reference sequence set. Sequence matching was performed in R version 4.5.2 using the Biostrings package. For each ZF probe, the target sequence was searched within the corresponding centromeric sequence (or sequences) using matchPattern. To account for different sequence orientation within the target region, the input sequence, reverse sequence, complement, and reverse-complement sequence were queried. Matches were then summarized as the number of target sites per probe and chromosome. To evaluate the presence of closely related target sequences, searches were performed allowing up to 6 mismatches between the query sequence and the reference. To account for potential centromeric off-target binding, the same search strategy was repeated across all centromeric sequences except the target chromosome for each probe. Matches detected on non-target centromeric sequences were counted as off-target centromeric hits and summarized per probe and off-target chromosome.

To assess the specificity of the ZF probes for different centromeres, we searched 2,110 complete and accurate centromeres assembled from 65 diverse human genomes^28^ for 224 ZF target sequences using a custom script we developed, findSeq.py. We found that almost all target sequences (222 out of 224) were present in at least one centromere within this dataset. We quantified the number of sites for each target sequence with a custom script, seqCount.py, and observed high concordance between the number of sites and the target chromosome (Figure 3a, S3a).

### Cell culture

HCT116 cells were grown in McCoy’s 5A (Modified) Medium supplemented with 10% FBS and Pen-Strep (Table S7). Media composition for *hTERT TP53⁻/⁻* human colonic epithelial cells (hCECs) is provided in Table S7 and in^19^. Cells were grown at 37C in a humidified 5% CO_2_ atmosphere.

### HCT116 cells were transfected using X-tremeGENE 360, following the manufacturer’s recommendations

For live microscopy, cells were grown on glass-bottom multi-well plates (#1.5H). For hCECs, glass wells were coated with 10 μg/ml fibronectin in PBS for 1h at 37C before cell plating. Prior to imaging, cells were incubated in a solution of 2-5μg/ml of Hoechst in culture medium for 10-15 min at 37C. When necessary, HaloTag was labeled by incubating in a 100nM solution of the appropriate Janelia Fluor (JF) dye in culture medium for 30 min, followed by x2 washes in 1X PBS (henceforth simply PBS) before replacing with culture medium and incubating for at least 25 min. Before imaging, Hoechst or JF media were replaced with FluoroBrite DMEM supplemented with 1X GlutaMAX, 10% FBS and 100U/ml Pen-Strep.

### Cell sorting

When necessary, Fluorescence-Activated Cell Sorting (FACS) was done using a Sony SH800 Cell Sorter set up in a laminar flow hood. Prior to sorting, cells were dissociated using TrypLE, diluted in culture medium, pelleted, resuspended in supplemented FluoroBrite DMEM to ∼10^6^ cells/ml and forced through a 35μm nylon strainer into 5ml round-bottom tubes, kept on ice.

### PiggyBac cell line generation

HCT116 lines expressing a single FP:ZF fusion were generated by transfection of equimolar amounts of the PiggyBac vector carrying the payload and a vector expressing the mammalian PiggyBac (mPB) transpose^26^ using XtremeGENE 360. 48h after transfection cells were treated with 1μg/ml puromycin for 5-7 days before release and expansion. No additional processing of the resulting polyclonal lines was done before imaging for ZF characterization.

Multi-color ZF lines were generated by transfecting cells with 2 or 3 PiggyBac vectors and an equimolar amount of the mPB vector and processed as above. After puromycin selection, the population was enriched in double or triple positive cells by FACS.

### Generation of mStayGold:dCas9 and mStayGold:Cen-ZF hCEC lines

An mStayGold:dCas9 hCEC line was generated by lentiviral infection of a diploid hCEC clone^19^, followed by antibiotic selection and fluorescence-activated cell sorting (FACS) (Figure S2a, b). Lentivirus were produced by transfecting HEK293T cells with the lentiviral pPAX and pMD2.G vectors, using Lipofectamine 3000. Medium was changed 12h after transfection. Virus-containing medium was collected 48 and 72h after transfection, filtered through a 0.45μm filter, mixed with Polybrene and added to hCEC cells. Cells were placed under appropriate antibiotic selection for 5 days starting 48h after the first infection.

To match the expression of dCas9 to that of the ZFs, another set of the same parental hCEC clone was infected with a lentiviral vector carrying an mStayGold:Cen7-ZF1 fusion and processed in parallel to the cells infected with mStayGold:dCas9 (Figure S2a). After drug selection, cells infected with mStayGold:dCas9 were sorted by FACS using a gate corresponding to the top 50% intensity of the mStayGold:Cen7-ZF1 population. This mStayGold:dCas9 hCEC population was expanded and used for all subsequent experiments. The nuclear fluorescence of these cells was found to be comparable to that of hCECs infected with other Cen-ZF in subsequent experiments (Figure S2d).

### dCas9 vs Cen-ZF comparison

For experiments comparing dCas9 and ZF labeling, ploidy and DNA damage, the parental hCEC clone was newly infected with mStayGold:Cen-ZFs at the same time that the mStayGold:dCas9 line was infected with lentivirus expressing the sgRNAs, and processed in parallel (Figure 2a, S2e). Cells were allowed to grow for 10 days post-infection, during which locus labeling was assessed. Similar nuclear fluorescence intensity and labeling efficiency were observed in all groups (Figure S2d-f). After 10 days in culture cell lines were frozen before further analysis.

### Targeted single-cell DNA sequencing and chromosome copy number profiling

hCEC cells infected with mStayGold:Cen-ZF lentivirus or mStayGold:dCas9 + sgRNAs were thawed and processed for targeted single-cell DNA sequencing using the Tapestri platform v3 kit, following the manufacturer’s protocol. Briefly, cells were washed in Ca^2+^- and Mg^2+^-free PBS, and filtered using 40μm cell strainers to obtain a single cell suspension. Uniquely barcoded samples of ∼200K cells per construct, with >80% viability, were pooled for a single Tapestri single-cell DNA sequencing run, spiking-in RPE1 cells as a diploid control. The final pool was resuspended in the provided cell buffer at a concentration of ∼4000 cells/µl, 35 µl of which was loaded onto the Tapestri DNA cartridge. Cell droplet encapsulation was performed in the Tapestri instrument with a CO-810 custom designed panel for the targeted single-cell DNA. The final targeted PCR amplicon library was sequenced using the Illumina NovaSeq X Plus on a 10B 300 cycles flowcell in a 2×150bp paired-end (PE) read format. The resulting sequencing data were processed using Tapestri pipeline DNA/standard version 3.6 for deconvolution of cell barcodes, read counts, and variant calling.

The Tapestri pipeline provides read counts per probe for each cell and variant allele frequencies for called variants for each cell. These data were further processed using the R package KaryoTapR^44^ to infer copy number profiles of individual cells at whole chromosome and arm levels.

### Aneuploidy induction

To induce aneuploidy, we treated cells with the MPS1 kinase inhibitor AZ3146. Polyclonal lines of HCT116 cells expressing multiple FP:Cen-ZF fusions were grown on sterilized, gelatinized 22×22mm 1.5H glass coverslips in 6-well dishes (see Immunofluorescence section, below). Cells were seeded sparsely (2 x 10^4^ cells/well) to allow clone formation. 24h after plating, cells were treated with either 5μM AZ3146 or with 0.1% DMSO as control. Treatment was maintained for 24h, followed by 48h in 0.1% DMSO, before fixing in 4% paraformaldehyde (PFA) for 10 min at RT, washed x3 in PBS, stained with 2μg/ml Hoechst in PBS for 5 min, washed in PBS and mounted by inverting coverslips onto 15-20μl drops of ProLong Gold antifade reagent on glass slides and allowing to cure in the dark for 24-48h at RT.

### Epigenetic editor recruitment

For RING1 recruitment to centromeres, HCT116 cells were seeded on glass coverslips, as described below (*Immunofluorescence*). Cells were transfected 24h later with the appropriate plasmids and treated with 1μg/ml Doxycycline (Dox) to induce the Tet-Responsive Element (TRE). The next morning, cells were treated with 5μM of Mandipropamid (in addition to Dox) — or treated with Dox only — for 6h to recruit the HaloTag:RING1:PYR1 fusion to the locus, after which cells were fixed in 4% PFA as described below. 1h prior to fixation, HaloTag was labeled with 100nM JFX650 for 30 min, before washing x2 with PBS and incubating in label-free medium for a further 25 minutes. Medium with HaloTag ligand also contained 1μg/ml Dox and 5μM Mandi. 5μM Mandipropamid was also added to 4% PFA to prevent release of PYR1 from ABI1 during the 10min fixation. Cells were subsequently immunolabeled as described below.

### Immunofluorescence and cell cycle analysis

Immunofluorescence was performed on cells growing on 22×22mm high precision (#1.5H) glass coverslips. Prior to cell plating, coverslips were sterilized by immersion in 70% ethanol for ∼30s and rinsed x2 with sterile, distilled water before coating (if needed). For HCT116 cells, coverslips were coated with 0.1% gelatin for 5 minutes at room temperature (RT) before cell plating. For hCECs, coverslips were covered with 5 μg/ml fibronectin in PBS for 1h at 37C before cell plating.

Cells were fixed in 4% PFA in PBS for 10 min at RT, washed x3 in PBS, permeabilized in 0.5% Triton X-100 for 10 min at RT, washed x2 in PBS and blocked for 30 min at RT in blocking buffer (2% Goat Serum, 2.5% bovine serum albumin (BSA), 0.1% Tween-20 in PBS)^45^. All primary antibodies were diluted in blocking buffer. Cells were incubated in primary antibodies for 1h at RT by inverting coverslips onto 30-50μl drops of primary antibody solution on sheets of Parafilm. Coverslips were subsequently washed x3 in wash buffer (0.2% BSA, 0.1% Tween-20 in PBS) before incubating in secondary antibodies for 1h at RT. Secondary antibodies were all diluted 1:500 in blocking buffer. Coverslips were subsequently washed x3 in 0.1% Tween-20 in PBS at RT before incubating in 2μg/ml Hoechst in PBS for 5 min at RT. Coverslips were washed in PBS for 5 min at RT before mounting. Samples were mounted by inverting coverslips onto 15-20μl drops of ProLong Gold antifade reagent on glass slides and allowed to cure in the dark for 24-48h at RT.

To evaluate cell cycle stage, cells were incubated in a 10μM solution of the thymidine analog EdU in culture medium for 1h prior to fixing, to label newly synthesized DNA. After immunofluorescence, and prior to staining DNA with Hoechst, coverslips were incubated for 30 minutes in the Click-iT^TM^ reaction cocktail (10% 1X Reaction Buffer Additive, 2% Copper protectant, 0.5% Fluorescent dye picolyl azide in PBS), as per manufacturer’s instructions (see Materials table), followed by a PBS wash, incubation in Hoechst, and mounting as described above.

### Halo-DNA FISH

For DNA FISH, cells were seeded and grown on coated glass coverslips, as described above for immunofluorescence. To avoid excessive cell detachment after pre-extracting soluble proteins, coverslips were coated with a solution of 10μg/ml Poly-D-Lysine for 15 minutes (instead of gelatin) and rinsed with PBS before plating cells.

Prior to DNA FISH, HaloTag was labeled with an appropriate JaneliaFluor® dye (JFX554, JFX650 or JF657, as necessary). The JF dye was diluted in culture medium to 100nM, cells incubated in the solution for 30 min, subsequently washed x2 with PBS and incubated again in fresh culture medium for at least 25 minutes before a final wash in PBS. To improve signal to background of HaloTag:ZF foci, soluble proteins were pre-extracted before fixing by incubating in cytoskeleton (CSK) buffer for 2 minutes^46^, and immediately fixed in freshly made 4% PFA in PBS for 10 min at room temperature (RT).

DNA FISH was performed in freshly fixed cells as described in{Clark.2026}. Briefly, cells were washed x3 in PBS after fixation, permeabilized for 10 min at RT in 0.5% Triton X-100, washed x2 in PBS, incubated for 5 min at RT in 0.1N HCl and washed x3 in 2X SSC and x3 in PBS. Cells were then incubated for 1h at 37C in a solution of 10μg/ml of RNAseA in PBS, washed x1 in PBS and incubated for 30 min at 37C in pre-hybridization buffer (2X SSC, 50% formamide, 0.1% Tween-20 in water). Centromeric FISH probes were either commercial (Table S6) or made in-house by direct labeling of amino-modified oligos (IDT) with AlexaFluor® 647 NHS ester. For hybridization, FISH probes were diluted in hybridization buffer (2X SSC, 50% formamide, 0.1% Tween-20, 10% dextran sulfate in water) at either 0.2 pmol/uL (home-made probes) or 0.025 nl/uL (OGT). Coverslips were inverted over ∼15-20uL of probe mix on glass slides, sealed with rubber cement and allowed to cure for 5-10min min the dark. Samples were denatured for precisely 5min at 85C by placing the slides on a thermalized metal block, after which slides were transferred to a humidified chamber and allowed to hybridize overnight at 42C. The next day, coverslips were returned to a multi-well dish and washed x2 in probe wash buffer (5X SSC, 50% formamide, 0.1% Tween-20 in water) for 30 min at 42C, x2 in 5X SSCT (5X SSC, 0.1% Tween-20 in water) and x1 in PBS before labeling DNA by incubating in a solution of 2μg/ml of Hoechst in PBS for 5min. After one final wash in PBS, cells were mounted by inverting coverslips onto 15-20μl drops of ProLong Gold antifade reagent on glass slides and allowed to cure in the dark for 24-48h at RT.

### Fluorescence imaging

Imaging was done using an epifluorescence Nikon Ti-E microscope fitted with lasers emitting at 405nm, 488nm, 532nm, 561nm, 594nm and 637nm and controlled by Micro-Manager. Samples were imaged through an oil-immersion Olympus 60X UPlanXApo (NA = 1.4) lens and images acquired with a Photometrics Prime BSI Express camera. Z-stacks of 2048 x 2048px images (effective pixel area of 98 x 98nm) were acquired every 250nm to capture the entire cell volume. All images in each experiment and its replicates were acquired with the same laser and exposure parameters.

### Image processing and analysis

All image analysis was done on maximum intensity projections (MIPs) of the Z-stacks to reduce computational burden. Multi-channel and single-channel MIPs for all images were generated using custom ImageJ macros^47,48^. Nuclei were detected using Cellpose 2.0^49^ on the MIP corresponding to the nuclear (Hoechst) staining for each image. Fluorescent foci were detected using Spotiflow^50^ on each of the relevant fluorescence channels for each image. Spotiflow was used to train a model to detect FISH foci (characterized by high brightness and very high signal to background ratio (SBR)), and a separate model to detect ZF foci (variable brightness and SBR). The FISH model was trained on FISH images and applied to all FISH images, as well as to IF images of proteins with similar pattern, such as CENP-B. The ZF model was trained on live still images of cells expressing a diversity of FP:ZF fusions, representative of the appearance of different ZF probes, and applied to all ZF fluorescence images, of either live or fixed cells.

We wrote custom Python code to do all image analysis. All images were processed as batches, defined by a single imaging session, with any normalization or calculations done separately on each batch. Each cell in each image and experiment is uniquely defined by an experiment ID + image ID + nuclear ID. Using multi-channel MIPs as input and masks generated by Cellpose to identify nuclei, we performed background subtraction on all nuclei (background minima) on a per-image basis, and outlier removal (nuclei with area outside mean ± 2*SD) on each batch before extracting basic statistics for each nucleus (cell) and each fluorescent channel in each image. Similarly, we used Cellpose nuclear masks and spots coordinates generated by Spotiflow to assign foci to each nucleus. Foci located outside of nuclei were discarded as false calls. Background subtraction was also performed on the spot intensity value calculated by Spotiflow for each image, and the SBR of each spot was calculated. To remove false positive foci, we filtered out spots with fluorescence intensity below 25% that of the brightest spot in each nucleus for each channel. No additional filtering steps were performed on fluorescent foci.

Spot co-localization was done by first calculating the 2D distance from each spot on a given fluorescence channel (query channel) to the nearest neighbor (NN) spot on one or more channels (target channels), and then assessing the distribution of NN distances. This distribution can be visualized as a cumulative distribution function (CDF) (Figure 1e) of the fraction of foci located within a given distance of their nearest neighbor. As a threshold for co-localization, we used 300nm (∼3 pixels in our images), which was the 2D distance between 95% of foci corresponding to the same locus (chr7 centromere) labeled with two FISH probes with the same DNA sequence but spectrally different fluorophores (median distance = 121nm) (Figure S1a, b). To determine ZF co-localization in Halo-FISH images, we used ZF foci as query and FISH foci as target channels, since all cells show FISH signal, but only a fraction had visible ZF foci.

Local fluorescence enrichment around a locus was calculated by defining a query channel (containing the locus of interest) and measuring the mean and total fluorescence intensity in the target channel(s) within a circle of 5px diameter (∼500nm) centered on the spot. Relative enrichment was calculated as the mean intensity of the target channel around the locus of interest divided by the corresponding nuclear median fluorescence intensity for each cell. The results of all analyses were stored as CSV files and plotted using R.

## Data and materials availability

All plasmids and cell lines are currently available upon request. Code for image analysis is available at https://github.com/timotheelionnet/ZATELLITE_image-analysis

## Acknowledgements

The authors would like to thank members of the Institute for Systems Genetics for feedback on this work, Albert Dominguez Mantes for guidance using Spotiflow. BioRender, licensed by NYU, was used for the creation of diagrams in figures 1, 2 and 5. This work was delivered as part of the eDyNAmiC team supported by the Cancer Grand Challenges partnership, funded in part by Cancer Research UK (J.D.B. CGCATF-2021/100018) and the National Cancer Institute (J.D.B. OT2CA278666) and also by NHGRI/NIH grant RM1-HG009491 to JDB. This work was supported by grants from the National Institutes of Health (R37 CA248631, R01 DK135089 and R01 R01 HG012590 to TD and R01GM149835 to TL) and grant PJT-190310 from the Canadian Institutes of Health Research to PMK. Compute resources from the Digital Research Alliance of Canada were used in this work.

## References

1. Misteli, T. The Self-Organizing Genome: Principles of Genome Architecture and Function. Cell 183, 28–45 (2020).

2. Bolzer, A. et al. Three-Dimensional Maps of All Chromosomes in Human Male Fibroblast Nuclei and Prometaphase Rosettes. Plos Biol 3, e157 (2005).

3. Beliveau, B. J. et al. Versatile design and synthesis platform for visualizing genomes with Oligopaint FISH probes. Proc National Acad Sci 109, 21301–21306 (2012).

4. Hsieh, T.-H. S. et al. Mapping Nucleosome Resolution Chromosome Folding in Yeast by Micro-C. Cell 162, 108–119 (2015).

5. Hua, P. et al. Defining genome architecture at base-pair resolution. Nature 595, 125– 129 (2021).

6. Takei, Y. et al. Spatial multi-omics reveals cell-type-specific nuclear compartments. Nature 641, 1037–1047 (2025).

7. Li, H. et al. Mapping chromatin structure at base-pair resolution unveils a unified model of cis-regulatory element interactions. Cell 188, 7175–7193.e19 (2025).

8. Robinett, C. C. et al. In vivo localization of DNA sequences and visualization of large-scale chromatin organization using lac operator/repressor recognition. J Cell Biology 135, 1685–1700 (1996).

9. Alexander, J. M. et al. Live-cell imaging reveals enhancer-dependent Sox2 transcription in the absence of enhancer proximity. Elife 8, e41769 (2019).

10. Gabriele, M. et al. Dynamics of CTCF- and cohesin-mediated chromatin looping revealed by live-cell imaging. Science 376, 496–501 (2022).

11. Lee, J. et al. Kinetic organization of the genome revealed by ultra-resolution, multiscale live imaging. (2025) doi:10.1101/2025.03.27.645817.

12. Chen, B. et al. Dynamic Imaging of Genomic Loci in Living Human Cells by an Optimized CRISPR/Cas System. Cell 155, 1479–1491 (2013).

13. Ma, H. et al. Multiplexed labeling of genomic loci with dCas9 and engineered sgRNAs using CRISPRainbow. Nat Biotechnol 34, 528–530 (2016).

14. Guan, J., Liu, H., Shi, X., Feng, S. & Huang, B. Tracking Multiple Genomic Elements Using Correlative CRISPR Imaging and Sequential DNA FISH. Biophys J 112, 1077– 1084 (2017).

15. Chen, B., Zou, W., Xu, H., Liang, Y. & Huang, B. Efficient labeling and imaging of protein-coding genes in living cells using CRISPR-Tag. Nat Commun 9, 5065 (2018).

16. Ma, H. et al. CRISPR-Sirius: RNA scaffolds for signal amplification in genome imaging. Nat. Methods 15, 928–931 (2018).

17. Clow, P. A. et al. CRISPR-mediated multiplexed live cell imaging of nonrepetitive genomic loci with one guide RNA per locus. Nat Commun 13, 1871 (2022).

18. Zhu, Y. et al. High-resolution dynamic imaging of chromatin DNA communication using Oligo-LiveFISH. Cell (2025) doi:10.1016/j.cell.2025.03.032.

19. Bosco, N. et al. KaryoCreate: A CRISPR-based technology to study chromosome-specific aneuploidy by targeting human centromeres. Cell 186, 1985–2001.e19 (2023).

20. Singh, M. et al. Elucidation of the molecular mechanism of the breakage-fusion-bridge (BFB) cycle using a CRISPR-dCas9 cellular model. Nucleic Acids Res. 52, 11689–11703 (2024).

21. Feng, H. et al. Genome-wide Chromosome-specific Aneuploidy Engineering and Phenotypic Characterization with CRISPR-Taiji. bioRxiv 2025.04.25.650684 (2025) doi:10.1101/2025.04.25.650684.

22. Ichikawa, D. M. et al. A universal deep-learning model for zinc finger design enables transcription factor reprogramming. Nat Biotechnol 1–13 (2023) doi:10.1038/s41587-022-01624-4.

23. Nurk, S. et al. The complete sequence of a human genome. Science 376, 44–53 (2022).

24. Altemose, N. et al. Complete genomic and epigenetic maps of human centromeres. Science 376, eabl4178 (2022).

25. Ivorra-Molla, E. et al. A monomeric StayGold fluorescent protein. Nat. Biotechnol. 42, 1368–1371 (2024).

26. Cadiñanos, J. & Bradley, A. Generation of an inducible and optimized piggyBac transposon system†. Nucleic Acids Res. 35, e87 (2007).

27. Miga, K. H. & Alexandrov, I. A. Variation and Evolution of Human Centromeres: A Field Guide and Perspective. Annu. Rev. Genet. 55, 583–602 (2021).

28. Gao, S. et al. A global view of human centromere variation and evolution. Nature 1– 12 (2026) doi:10.1038/s41586-026-10841-9.

29. Ma, H. et al. Multicolor CRISPR labeling of chromosomal loci in human cells. Proc. Natl. Acad. Sci. 112, 3002–3007 (2015).

30. Ben-David, U. & Amon, A. Context is everything: aneuploidy in cancer. Nat. Rev. Genet. 21, 44–62 (2020).

31. Hewitt, L. et al. Sustained Mps1 activity is required in mitosis to recruit O-Mad2 to the Mad1–C-Mad2 core complex. J. Cell Biol. 190, 25–34 (2010).

32. Ziegler, M. J. et al. Mandipropamid as a chemical inducer of proximity for in vivo applications. Nat. Chem. Biol. 18, 64–69 (2022).

33. Xu, X. et al. Engineered miniature CRISPR-Cas system for mammalian genome regulation and editing. Mol. Cell 81, 4333–4345.e4 (2021).

34. Ma, H., Reyes-Gutierrez, P. & Pederson, T. Visualization of repetitive DNA sequences in human chromosomes with transcription activator-like effectors. Proc. Natl. Acad. Sci. 110, 21048–21053 (2013).

35. Thanisch, K. et al. Targeting and tracing of specific DNA sequences with dTALEs in living cells. Nucleic Acids Res. 42, e38–e38 (2014).

36. Ma, H. et al. CRISPR-Cas9 nuclear dynamics and target recognition in living cells. J. Cell Biol. 214, 529–537 (2016).

37. Khalil, A. S. et al. A Synthetic Biology Framework for Programming Eukaryotic Transcription Functions. Cell 150, 647–658 (2012).

38. Halstead, J. M. et al. An RNA biosensor for imaging the first round of translation from single cells to living animals. Science 347, 1367–1671 (2015).

39. Bergmann, J. H. et al. Epigenetic engineering shows H3K4me2 is required for HJURP targeting and CENP-A assembly on a synthetic human kinetochore. EMBO J. 30, 328–340 (2011).

40. Martins, N. M. C. et al. Epigenetic engineering shows that a human centromere resists silencing mediated by H3K27me3/K9me3. Mol. Biol. Cell 27, 177–196 (2016).

41. Salinas-Luypaert, C. et al. DNA methylation influences human centromere positioning and function. Nat. Genet. 57, 2509–2521 (2025).

42. Los, G. V. et al. HaloTag: A Novel Protein Labeling Technology for Cell Imaging and Protein Analysis. ACS Chem. Biol. 3, 373–382 (2008).

43. Oren, Y. et al. Cycling cancer persister cells arise from lineages with distinct programs. Nature 596, 576–582 (2021).

44. Mays, J. C., et al. KaryoTap Enables Cost-Effective High-Throughput Single-Cell Aneuploidy Profiling and Reveals How Positive and Negative Selection Shapes Cancer Genomes. bioRxiv 2023.09.08.555746 (2025) doi:10.1101/2023.09.08.555746.

45. Chaumeil, J., Micsinai, M. & Skok, J. A. Combined Immunofluorescence and DNA FISH on 3D-preserved Interphase Nuclei to Study Changes in 3D Nuclear Organization. J. Vis. Exp. e50087 (2013) doi:10.3791/50087.

46. Whelan, D. R. & Rothenberg, E. Homologous Recombination, Methods and Protocols. Methods Mol. Biol. 2153, 355–363 (2020).

47. Schindelin, J., et al. Fiji: an open-source platform for biological-image analysis. Nat. Methods 9, 676–682 (2012).

48. Rueden, C. T. et al. ImageJ2: ImageJ for the next generation of scientific image data. BMC Bioinform. 18, 529 (2017).

49. Pachitariu, M. & Stringer, C. Cellpose 2.0: how to train your own model. Nat Methods 19, 1634–1641 (2022).

50. Mantes, A. D. et al. Spotiflow: accurate and efficient spot detection for fluorescence microscopy with deep stereographic flow regression. Nat. Methods 22, 1495–1504 (2025).

51. Xiong, H. et al. A highly stable monomeric red fluorescent protein for advanced microscopy. Nat. Methods 22, 1288–1298 (2025).

