## Supplementary Figures for "*ZATELLITE:* a toolkit to visualize and manipulate human centromeres in live cells using synthetic zinc fingers"

#### Figure 1, supplement

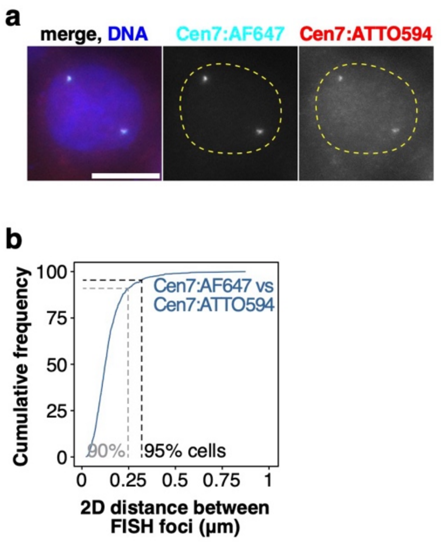


**Figure 1, supplement.**
**a.** Representative images of DNA FISH on HCT116 cells as a reference for co-localization. Cells were fixed and labeled with two FISH probes targeting the same DNA sequence in the centromere of chr7, but tagged with two different fluorophores, as indicated. **b.** Cumulative frequency distribution of 2D distances between both FISH foci in a population of cells like those in a. 95% of foci are within 300nm of their nearest neighbor (black dashed line); 90% of foci are within 250nm of their nearest neighbor (gray dashed line). Scale bars = 10µm.

### Figure 2, supplement 1

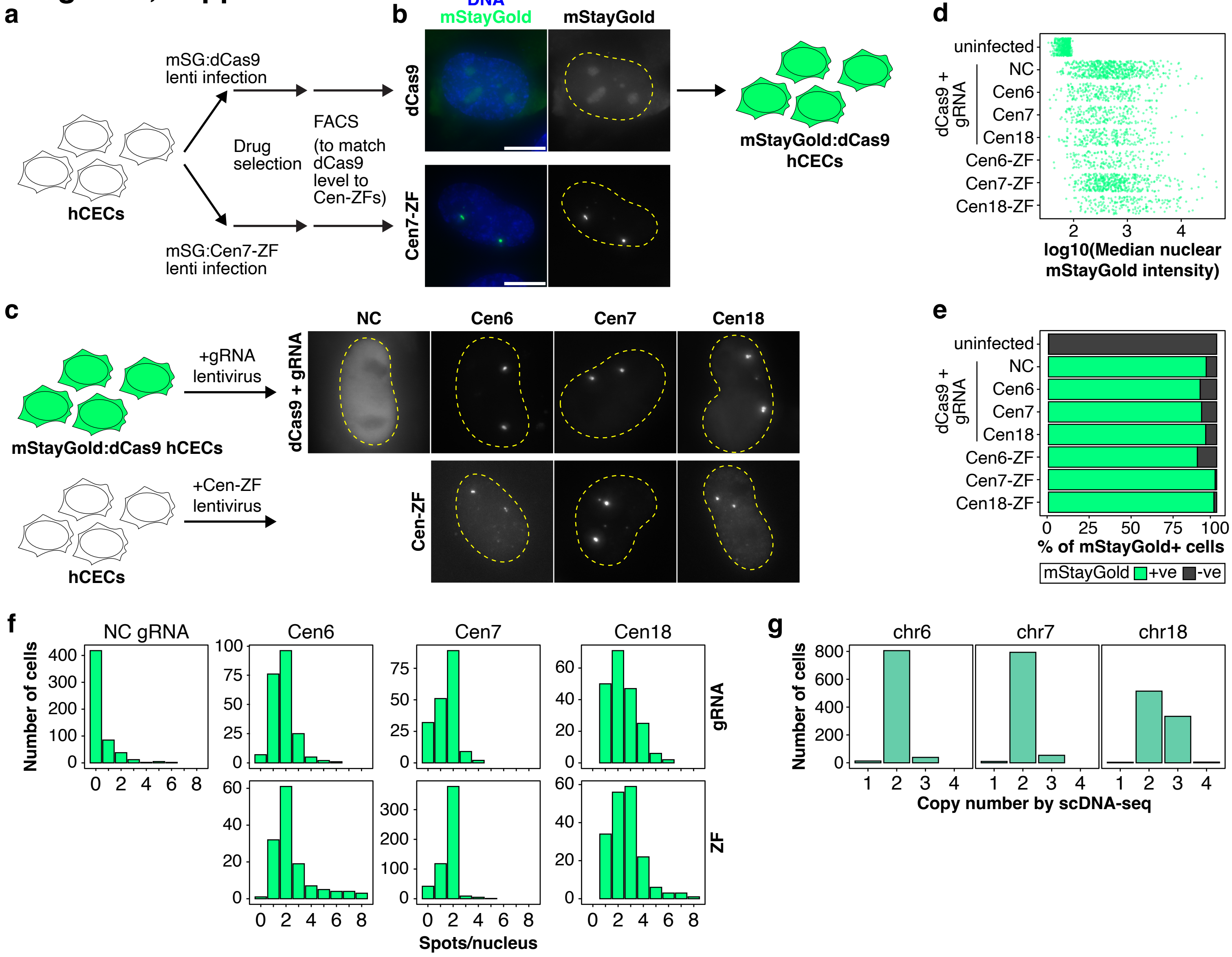

### Figure 2, supplement 2

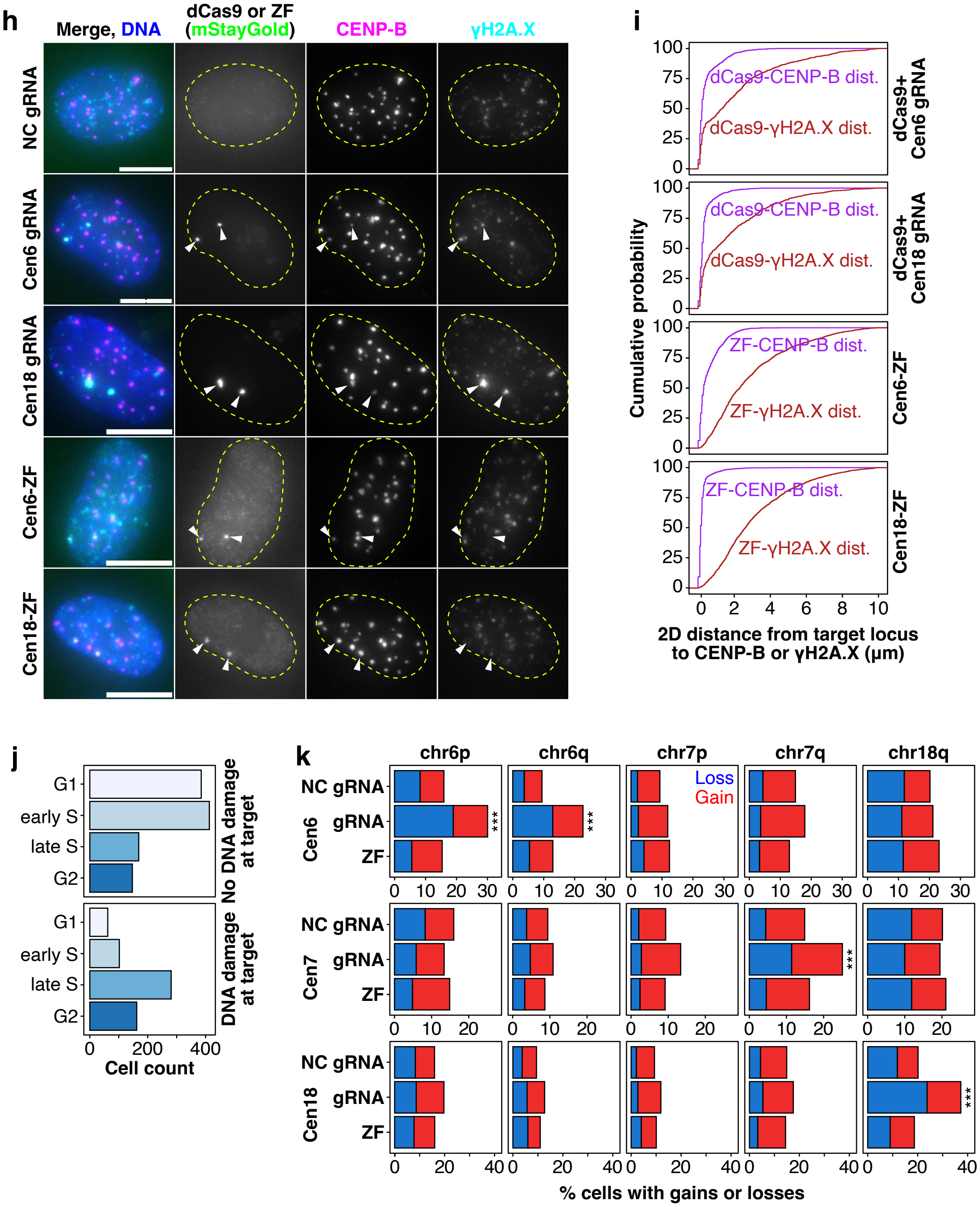

**Figure 2, supplement.**

**a**. Experimental diagram of mStayGold:dCas9 hCEC generation. hCECs were infected with a lentivirus producing an mStayGold:dCas9 fusion, and, in parallel, with lentiviruses producing mStayGold:Cen7-ZF fusions. mStayGold:dCas9 cells were sorted by FACS to match nuclear fluorescence levels to those of the mStayGold:Cen7-ZF fusions. **b**. Representative images of cells expressing mStayGold:dCas9 or mStayGold:Cen7-ZF1 after FACS sorting. **c**. Experimental diagram and representative images for experiment in Figure 2a showing images of hCECs with cen6, cen7 and cen18 labeled by each of the two labeling modalities. mStayGold:dCas9 cells generated in (a-c) were infected with sgRNAs. Parental hCECs were re-infected with mStayGold:Cen7-ZFs. **d**. Median nuclear mStayGold intensity in each of the groups shown in c. Each dot represents one nucleus. **e**. Fraction of mStayGold+ cells in each of the groups shown in c-d, as determined by automatic nuclear segmentation and classification (see Methods). **f**. Histograms showing the distribution of the number of fluorescent foci/nucleus in each of the experimental groups shown in (c). Fluorescent foci were counted automatically and filtered according to their signal to background ratio, as described in the Methods. **g**. Histograms showing the copy number for chr6, 7 and 18 in the parental hCEC line as determined by scDNA-seq. **h**. Representative images of hCECs from the same population as in c, fixed 10 days post-infection and stained for γH2A.X and CENP-B, as in Figure 2b. Arrowheads indicate the position of the mStayGold (dCas9-sgRNA or ZF) foci across channels. **i**. Cumulative frequency distributions of the 2D nearest-neighbor (NN) distance between mStayGold foci and centromeres (marked by CENP-B, magenta) or γH2A.X foci (dark red). Distances below 300nm indicate co-localization. **j**. Cell cycle distribution of cells without or with γH2A.X foci at the target locus. **k**. Bar plots showing the fraction of cells with gains or losses of the chromosomal arm indicated (top) in each of the experimental groups (non-coding (NC) gRNA, and gRNAs or ZFs targeting the indicated centromeres), as determined by scDNA-seq. The parental line was found to have an additional copy of chr18p (panel g), so gains/losses for that arm could not be assessed and is therefore not shown. *** = $p<0.001$ (Fisher's Exact test). Scale bars = 10µm.

### Figure 3, supplement

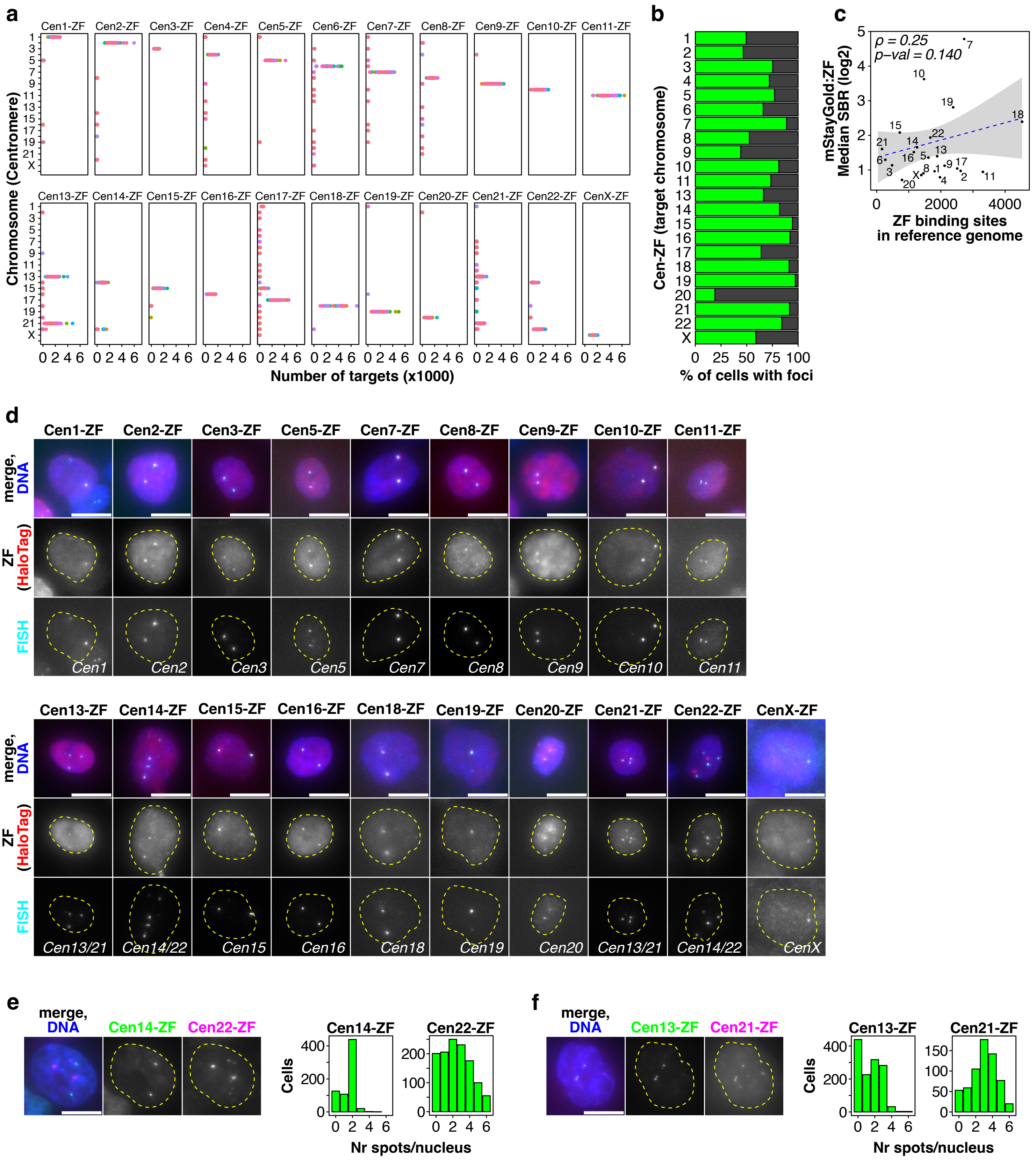

**Figure 3, supplement.**

**a**. Dot plot indicating the number of targets found for each of the Cen-ZF probes in each of 2110 human centromeres from a cohort of 65 individuals (color-coded)[28], indicating target specificity for each probe. **b**. Fraction of mStayGold+ cells showing fluorescent foci for each of the probes shown in Figure 3b (green: fraction of cells with foci, dark gray: fraction of cells without foci). **c**. Scatter plot showing the relationship between the number of binding sites on the T2T reference genome for each Cen-ZF and its signal to background ratio in live images (expressed as log2). Rho and p-value are shown (Spearman rank-order correlation test). **d**. Representative images of HCT116 cells stably expressing HaloTag:ZF fusions of the probes indicated, and subject to DNA FISH for their target locus, as in Figure 3c. HaloTag, labeled with JFX554, indicates the location of the *ZATELLITE* probe. The target foci for each probe (FISH) are marked by fluorescently labeled oligonucleotides. **e**. Representative images of live HCT116 cells stably expressing ZATELLITE fusions labeling both Cen14 (mStayGold) and Cen22 (HaloTag) and histograms showing the number of foci per nucleus for each probe in a representative sample of live cells for each Cen-ZF. The two bright, non-diffraction limited blobs for Cen22-ZF are likely the result of off-target binding (see validation in panel d). **f.** Same as (e) for Cen13-ZF (mStayGold) and Cen21-ZF (HaloTag). Scale bars = 10μm.
